# Perception of RNA Nanotechnology among students and recommendations towards improved educational outreach

**DOI:** 10.1101/2022.02.06.479286

**Authors:** Seraphim Kozlov

## Abstract

Translating new technologies to industrial and biomedical applications requires a highly skilled workforce. In the past, colleges and graduate schools played a primary role in preparing students for various areas of industry and medicine. The learning process and introduction of new concepts have recently extended beyond college education. High schools saw the rise of specialized career programs, while both high and middle school curricula got infused with challenging concepts. Nucleic acids, such as DNA and RNA, are broadly known for their role in the foundation of life. However, nucleic acid nanotechnology, an area of material science manipulating DNA and RNA to create complex structures with controlled properties, is less known. Herein, I report the results of a study investigating the perception of RNA nanotechnology among school students and suggest educational resources to improve understanding of RNA nanotechnology.

## Introduction

RNA Nanotechnology is a new area of science. It manipulates RNA oligonucleotides to assemble them into three-dimensional structures with different sizes, shapes, and functionalities (1, 2). The synthesis of RNA nanoparticles and the proof of principle of their applications in biology and medicine including cancer therapy, gene therapy, and vaccines are well established (2-4). Academic leaders of RNA nanotechnology formed a professional society, the International Society of RNA Nanotechnology and Nanomedicine to “*promote advances in basic sciences, medicine, pharmaceutical sciences, imaging, diagnostics, and various nanobiotechnological applications*.” (https://www.isrnn.org/). The Society is currently led by Professor Peixuan Guo of the Ohio State University.

While RNA is known to the public for its role in nature and live organisms, the knowledge of RNA Nanotechnology is mainly limited to the university labs creating these novel materials. As RNA Nanotechnology continues to grow, researchers are discussing the commercialization and translation of these novel materials to industrial and biomedical applications (2, 3).

Advancing new therapies from the bench to the clinic involves pre-clinical studies and clinical trials, and collaboration between academic researchers, medical doctors, and pharmaceutical scientists (5). On average, it takes about 25 to 30 years for a newly discovered drug to reach commercial use; the more complex a new technology, the longer this process takes (5). The importance of improving graduate students’ education in various disciplines involved in nanomedicine has recently been raised by Barton et al. (6).

Translating a sophisticated technology, such as RNA Nanotechnology, to the clinic also requires a well-educated workforce. Education in this area of science would benefit from starting earlier than college in one’s academic path. Therefore, to help educators and researchers in the field of RNA Nanotechnology to understand the perception of this field by students and to prepare helpful educational resources, I conducted my study.

Herein, I report the results of a survey probing undergraduate (middle school, high school, and college) students about their views on RNA Nanotechnology and the types of educational resources they want to have to advance their education in this area of science.

## Methods

A survey contained 12 questions, 11 of which were mandatory and 1 was optional; 4 questions allowed multiple answers (**Table 1**). The survey was announced on the webpage of the Biomedical and Life Sciences Youth Society https://www.blyseducation.org/. It was also populated via LinkedIn and through the author’s contacts. The survey remained open from July 10 to December 1, 2021. The identity of respondents such as name and affiliation was not tracked to respect their privacy and in compliance with 20 CFR 431.104 (d)(2)(i) and 20 CFR 431.104 (d)(2)(ii).

**Table 1.** Survey Questions. Details about questions and answers are summarized in the table. N/A = not applicable

| Number | The answer is required, Yes/No | Multiple answers are allowed, Yes/No | Question | Answer Options |
| --- | --- | --- | --- | --- |
| 1 | Yes | No | Which of the following best describes your school? | a. Middle school<br>b. High school<br>c. Undergraduate/College<br>d. Graduate<br>e. I am not a student |
| 2 | Yes | No | Please share your geographic location | a. US<br>b. UK<br>c. Brazil<br>d. Hong Kong<br>e. Hungary<br>f. Other (name) |
| 3 | Yes | N/A | What state in the US are you residing in? If not applicable, please type N/A | Free response |
| 4 | Yes | No | Who determines the types of classes you take at the school? | a. You<br>b. School<br>c. Parents/Guardians<br>d. None of the above |
| 5 | Yes | Yes | Which of the following classes have you taken in the past or will be taking in the upcoming school year? Please select all that apply | a. Biology<br>b. AP Biology<br>c. Chemistry<br>d. AP Chemistry<br>e. Biochemistry<br>f. Cell Biology<br>g. None of the above |
| 6 | Yes | No | Did you ever participate or do you plan on participating in a science fair competition? | a. Yes<br>b. No<br>c. Maybe |
| 7 | Yes | Yes | Have you ever presented your research at a scientific conference? | a. Yes, I presented a poster<br>b. Yes, I presented a speech<br>c. No |
| 8 | Yes | No | Do you know what RNA is? | a. Yes<br>b. No |
| 9 | Yes | Yes | What aspects of RNA do you find the most challenging? Please select all that apply | a. Complementarity<br>b. Its secondary structure<br>c. Its difference from DNA<br>d. Thermodynamics<br>e. Other |
| 10 | Yes | No | Are you aware of RNA Nanotechnology? | a. Yes<br>b. No |
| 11 | Yes | Yes | What resources would you like to have to learn more about RNA nanotechnology? Please select all that apply | a. Webinars<br>b. Hands-on activities<br>c. Mentor conducting research in this field<br>d. Internship in a research institution<br>e. Volunteering opportunity in a research lab |
| **12** | No | No | Please share your suggestions for what can be done to improve the quality of education in this field and increase awareness of RNA nanotechnology | Free response |

## Results and Discussion

The survey received seventy-one responses. Most respondents (56%) were from the US; the remaining 44% were from other countries as follows: India (14%); Brazil (6%); UK (4%); Chile (3%); Denmark (3%); United Arab Emirates (3%); Malaysia (3%); Iran (2%); Israel (2%); South Korea (1%); Spain (1%); Nigeria (1%); Sri Lanka (1%) (**Figure 1A**). Most of the US participants (68%) were from Maryland; the remaining 32% were from other states as follows: Colorado (7%); New York (5%), North Carolina (5%); Minnesota (3%); South Carolina (3%); Utah (3%); California (2%); Illinois (2%) and Massachusetts (2%) (**Figure 1A**).

**Figure 1.**
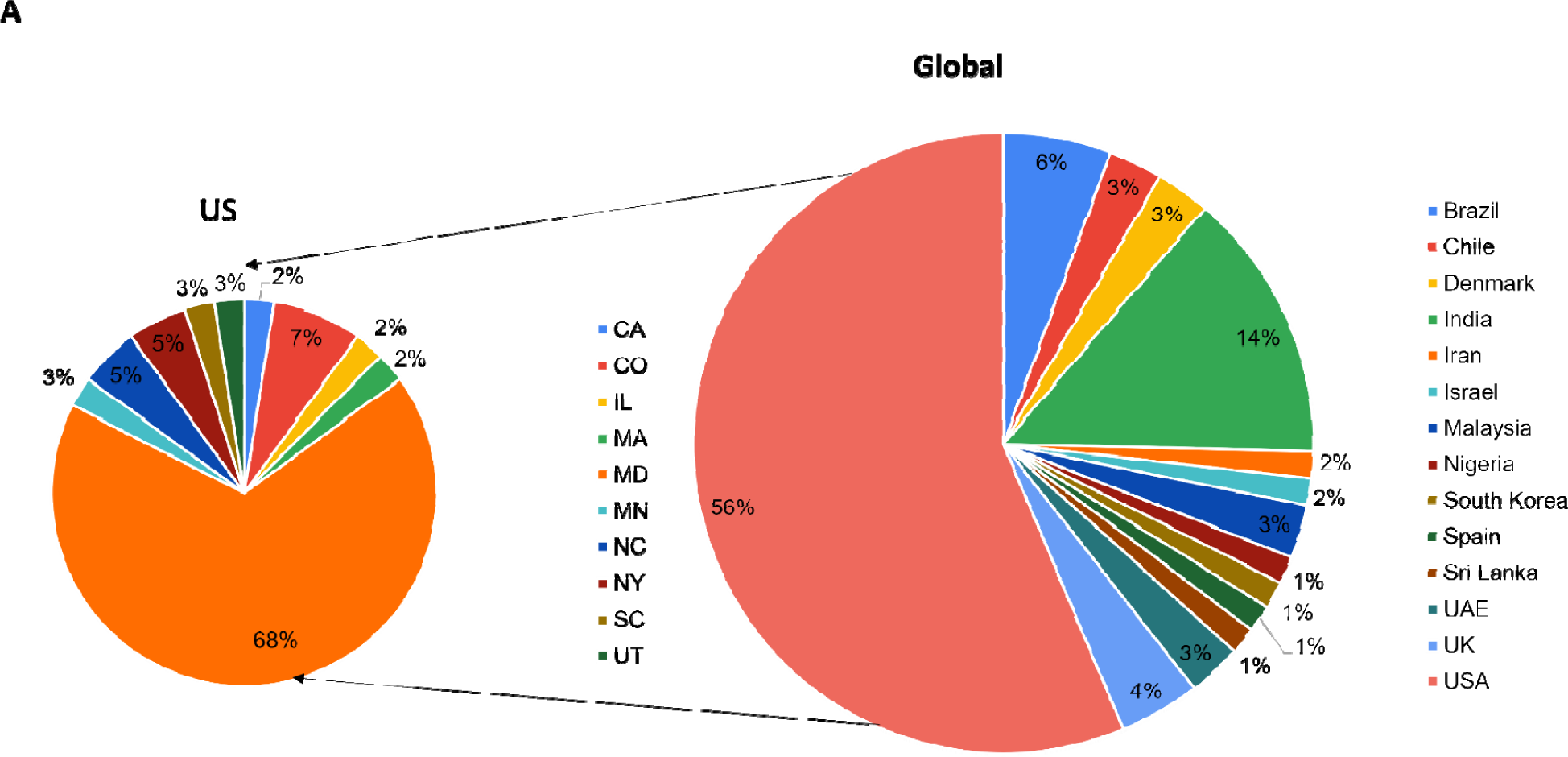
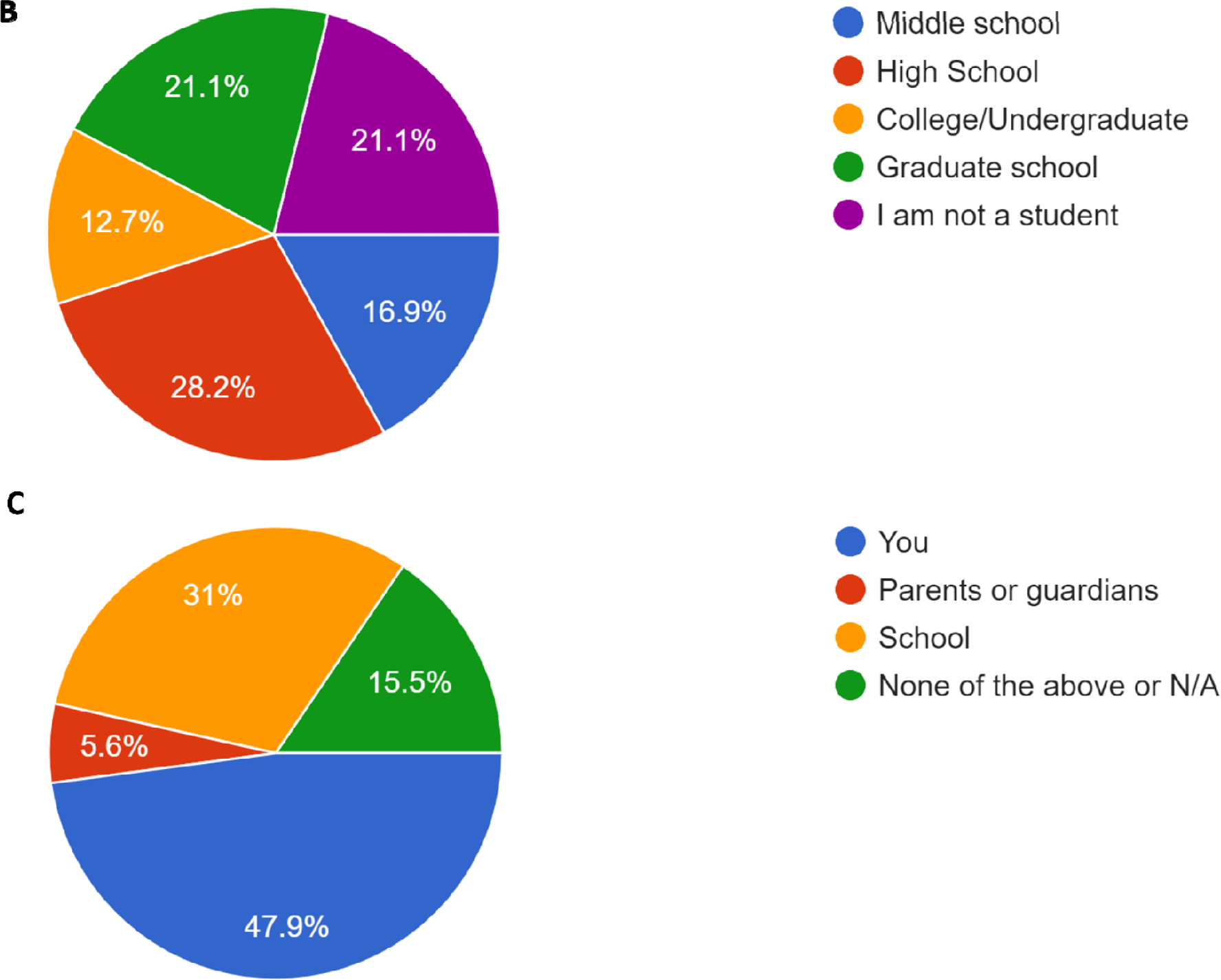
Participants’ demographic. Geographic location (A); type of school (B); who decides types of classes a student takes (C). Abbreviations: UAE = United Arab Emirates; UK = United Kingdom; USA = United States of America; CA = California; CO = Colorado; IL = Illinois; MA = Massachusetts; MD = Maryland; MN = Minnesota; NC = North Carolina; NY = New York; SC = South Carolina; UT = Utah; N/A = not applicable

High school students dominated among the respondents (28.2%). The breakdown of other respondents was as follows: graduate students (21.1%), adults (21%), middle school students (16.9%), and college undergraduate students (12.7%) (**Figure 1B**). In most cases (47.9%) students had the freedom of selecting the types of classes they take at school; in 31% of cases, schools determined that decision; parents or guardians had little influence (5.6%) (**Figure 1C**).

Biology and Chemistry were the most popular classes taken by respondents (54.9 and 49.3%, respectively). Other advanced classes including AP Biology (21.1%), AP Chemistry (29.6%), Biochemistry (29.6%), and Cell Biology (28.2%) were also taken. Twenty-two and a half percent of respondents (22.5%) did not take the above-mentioned classes (**Figure 2**). These respondents were middle school students, whose curriculum does not include these disciplines.

**Figure 2.**
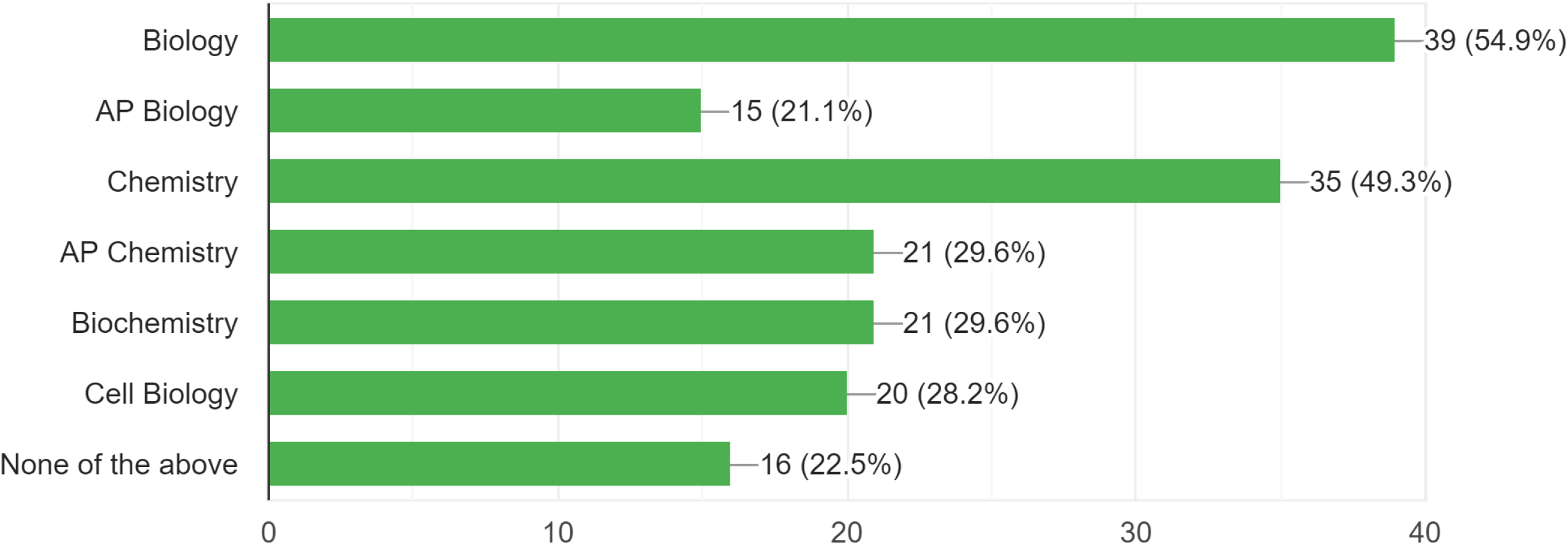
Types of classes taken by students before the survey. Participants were allowed to choose more than one response

The majority of respondents (53%) had a record of participation in a Science Fair event; 37% reported no participation and 10 % considered potentially participating in a Science Fair event (**Figure 3A**). The breakdown for participation in research conferences was as follows: 30% presented a poster, 32% gave a speech, and 38% did not participate in a conference (**Figure 3B**). Most respondents who did not participate in a research conference were from either high or middle school.

**Figure 3.**
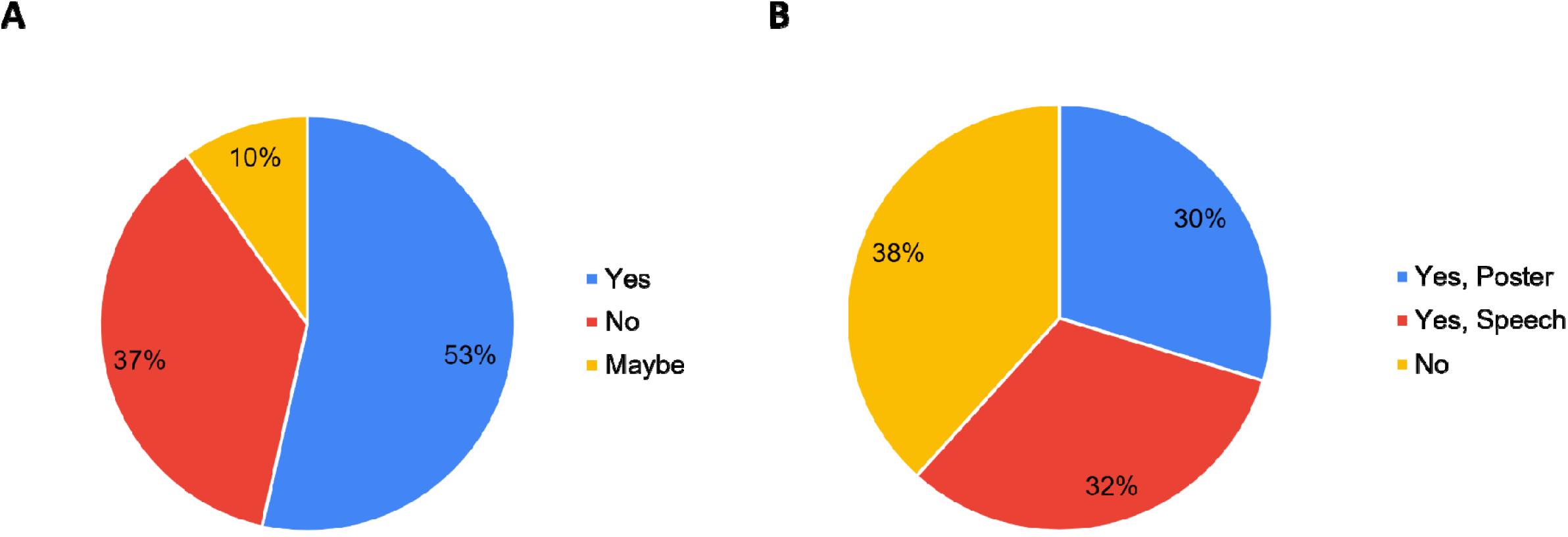
Participation in science-related events. Science Fair (A), Research conference (B). Respondents were allowed to choose more than one answer on the graph shown in B.

The awareness of RNA was very high, 91.5% (**Figure 4A**); the low percentage of respondents (8.5%) unaware of RNA were likely rising 6^th^ graders, who would learn about RNA during Science classes in 6^th^ grade. Despite high awareness of RNA, more than half of respondents (54.9%) were unaware of RNA nanotechnology; the remaining 45.1% of participants who responded positively to this question were graduate students and adults with a small proportion of undergraduate students (**Figure 4B**).

**Figure 4.**
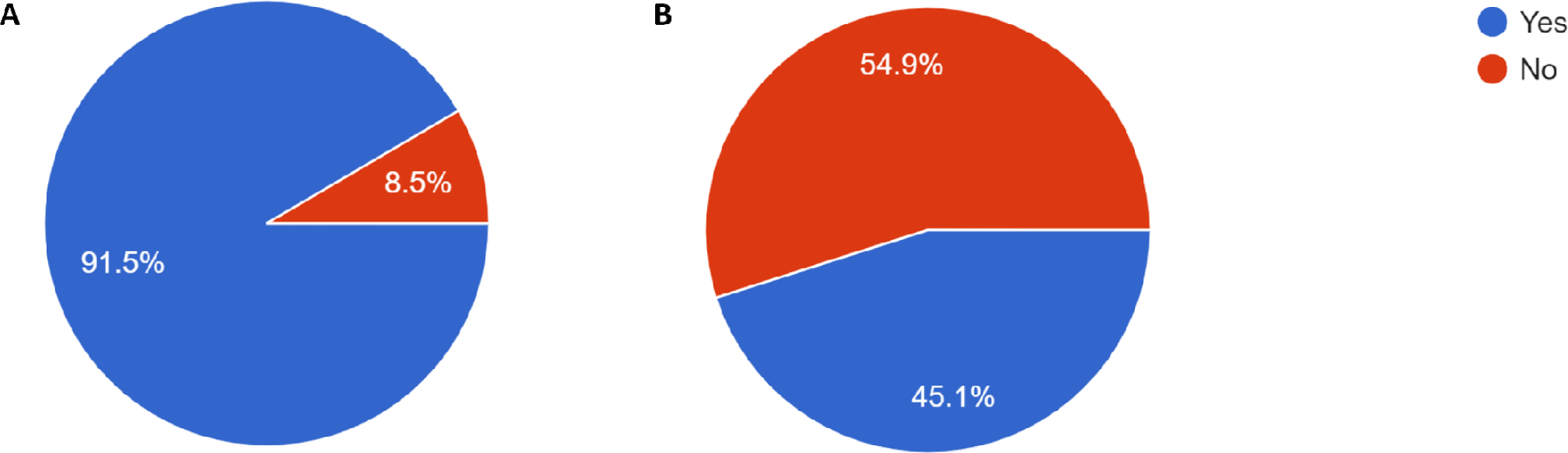
Awareness of RNA vs. RNA Nanotechnology. Awareness of RNA (A) and RNA Nanotechnology (B)

RNA secondary structure and thermodynamics were the most challenging concepts for most students, 57.7% and 64.8% of respondents, respectively (**Figure 5A**). Most respondents preferred to learn about RNA nanotechnology via webinars (70.4%), internship in a research lab (53.5%), hands-on activities (52.1%), and volunteering opportunities at research organizations (50.7%) (**Figure 5B**). Other ideas for educational activities that are needed to further improve education in the area of RNA nanotechnology as suggested by the survey respondents are summarized in **Table 2**.

**Table 2.** Respondents’ suggestions for improving education in the field of RNA Nanotechnology.

|  |
| --- |
| Provide training to teachers so that they can teach this information during regular classes |
| Network as cooperative integration with other educational nanotechnology initiatives as NaQaa Nanotechnology Network (NNN) www.nakaanetwork.webs.com |
| Increase awareness of its relevance to health and industry |
| Organize internships under the mentorship of eminent professors |
| Provide free access to education materials |
| Conduct webinars and hands-on training to improve the quality of education |
| Increase the visibility of the marketed and under-development agents that involve RNA nanotechnology |
| Raise the quality of education in general |
| Consider nanoformulation |
| Make it more interesting to students |
| Provide digital view and experiment |
| Learn it during science class |
| Create educational movies |
| Organize a school club |
| Create YouTube video |
| Given the current use of RNA based vaccines in combating Covid 19, more focus needs to be driven towards nano-based |
| techniques in increasing the efficacy of such vaccines for other diseases also |
| Provide more webinars on basics and current research on RNA nanotechnology |
| Incentify taking biology at a higher level |
| Increase awareness of the RNA use in vaccines |
| Promote this area at a higher level |
| Organize science fair or other educational events at which students could meet researchers investigating RNA |

**Figure 5.**
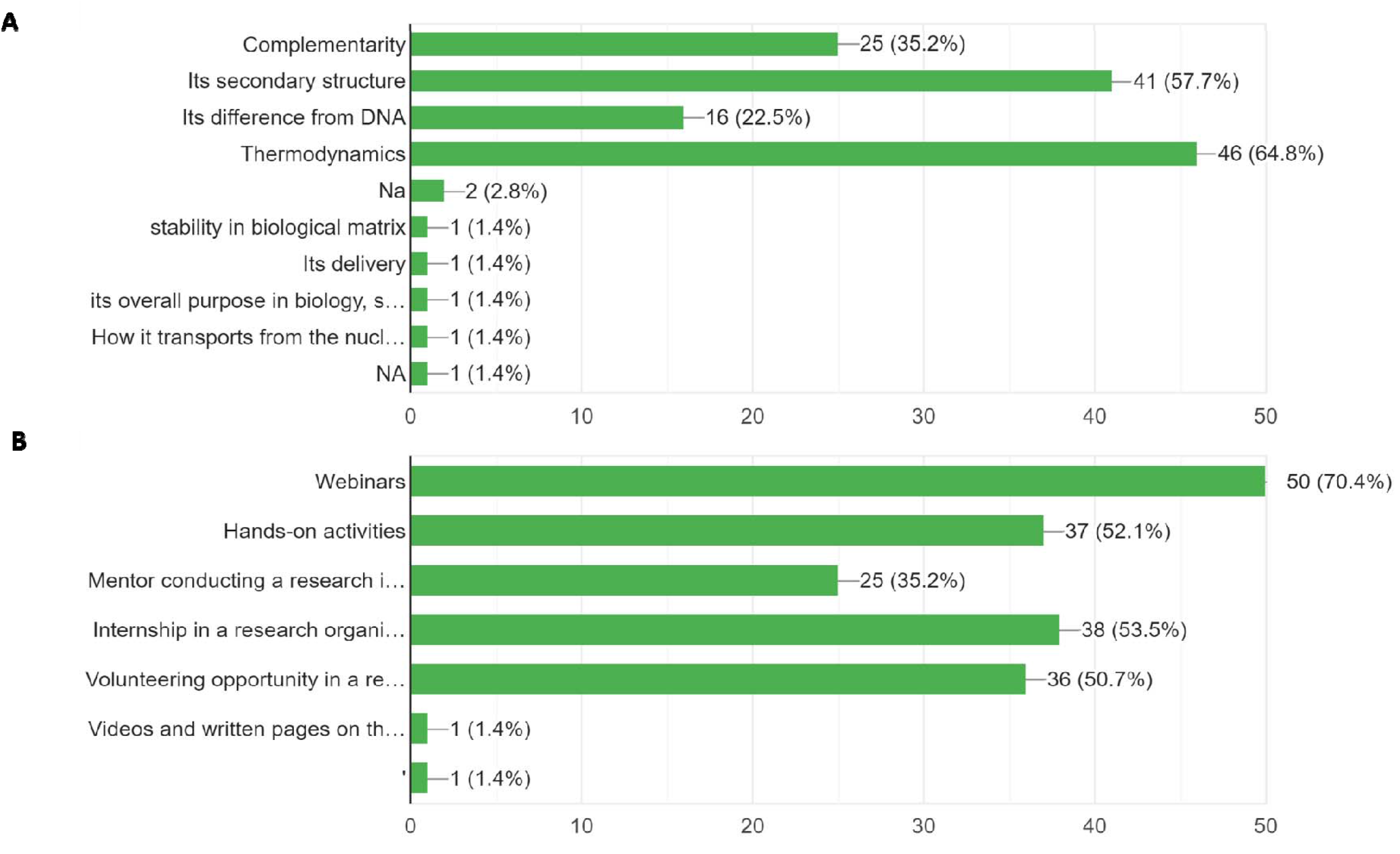
Answers regarding challenging areas of RNA and resources needed to improve the education. What aspects of RNA do you find most challenging? (A) What resources would you like to have to learn more about RNA Nanotechnology? (B). More details about questions and answer choices are provided in Table 1. Na = not applicable; s = science; nucl. = nucleus;

## Conclusions

The awareness of RNA by far exceeds that of RNA Nanotechnology. The majority of middle and high school students, as well as college undergraduate students, would benefit from educational activities including webinars, educational videos, hands-on activities, volunteering, and internships in an organization conducting active research in the RNA Nanotechnology field. International collaborations may further improve the outcome.

This report summarizes the opinion of a small cohort of students representing various countries. A study involving a larger number of students is needed to allow a quantitative comparison of results.

## Ethics

In the course of this project, the information was obtained by the investigator in such a manner that the identity of the human subjects cannot readily be ascertained, directly or through identifiers linked to the subjects. The disclosure of the study results would not reasonably place survey participants at risk of criminal or civil liability or be damaging to their financial standing, employability, educational advancement, or reputation. This study is, therefore, exempt from the IRB review process according to the 20 CFR 431.104 (d)(2)(i) and 20 CFR 431.104 (d)(2)(ii).

## Acknowledgments

I am grateful to Dr. Afonin for introducing me to RNA Nanotechnology, for his guidance and inspiration. I am also grateful to all survey participants for sharing their opinion. Provide training to teachers so that they can teach this information during regular classes

## References

1. Kim J, Franco E. RNA nanotechnology in synthetic biology. Curr Opin Biotechnol. 2020 Jun;63:135–141. doi: 10.1016/j.copbio.2019.12.016. Epub 2020 Feb 5. PMID: 32035339

2. Jasinski D, Haque F, Binzel DW, Guo P. Advancement of the Emerging Field of RNA Nanotechnology. ACS Nano. 2017 Feb 28;11(2):1142–1164. doi: 0.1021/acsnano.6b05737. Epub 2017 Feb 7. PMID: 28045501; PMCID: PMC5333189.

3. Johnson MB, Chandler M, Afonin KA. Nucleic acid nanoparticles (NANPs) as molecular tools to direct desirable and avoid undesirable immunological effects. Adv Drug Deliv Rev. 2021 Jun;173:427–438. doi: 10.1016/j.addr.2021.04.011. Epub 2021 Apr 20. PMID: 33857556; PMCID: PMC8178219.

4. Dao BN, Viard M, Martins AN, Kasprzak WK, Shapiro BA, Afonin KA. Triggering RNAi with multifunctional RNA nanoparticles and their delivery. DNA RNA Nanotechnol. 2015 Jan;2(1):1–12. doi: 10.1515/rnan-2015-0001. Epub 2015 Jul 27. PMID: 34322586; PMCID: PMC8315566.

5. Goldblatt EM, Lee WH. From bench to bedside: the growing use of translational research in cancer medicine. Am J Transl Res. 2010 Jan 1;2(1):1-18. PMID: 20182579; PMCID: PMC2826819.

6. Barton AE, Borchard G, Wacker MG, Pastorin G, Saleem IY, Chaudary S, Elbayoumi T, Zhao Z, Flühmann B. Need for Expansion of Pharmacy Education Globally for the Growing Field of Nanomedicine. Pharmacy. 2022; 10(1):17. https://doi.org/10.3390/pharmacy10010017

